# Integrating physiology and transcriptomics reveals the mechanisms underlying the differences in cotton (*Gossypium hirsutum* L.) growth and development under different photoperiods

**DOI:** 10.1101/2025.11.21.689390

**Authors:** Ning Zhang, Yujie Liu, Haitao Dai, Penghui Yi, Weiwei Cheng, Xinyu Zhan, Aiyu Liu, Xiaoju Tu

## Abstract

Photoperiod is a crucial environmental factor that regulates plant growth and flowering. While the photoperiodic flowering pathway has been extensively studied, the mechanisms underlying photoperiodic regulation of cotton (*Gossypium hirsutum* L.) growth and development remain unclear. In this study, we selected different genotypes of cotton varieties, XJ12 and XJ21-11, to investigate the effects of long-day (LD) and short-day (SD) conditions on cotton photosynthesis, dry matter accumulation, flowering time, and related gene expression, utilizing physiological index measurements, transcriptomics, and WGCNA analysis. The results indicated that LD treatment significantly enhanced photosynthetic parameters and dry matter weight in both cotton varieties, with the net photosynthetic rate (Pn) being 154.79% and 111.86% higher under LD compared to SD treatment. Additionally, the flowering time of the two varieties was advanced by 12 and 13 days, respectively, under LD treatment relative to SD treatment. LD conditions promoted gene expression in sucrose and starch metabolism, facilitating cotton growth. Furthermore, LD inhibited the expression of genes associated with flavonoid biosynthesis, thereby altering the secondary metabolism of cotton, and activated transcription factors through signaling pathways (including phytohormone signaling and MAPK signaling pathways) and biological clock rhythms, which induced earlier flowering in cotton. This study provides new insights into the molecular mechanism of the effect of photoperiod on cotton growth and development.

## 1. Introduction

Cotton is the world’s most significant cash crop, serving as a primary raw material for the textile industry, and is cultivated in over 75 countries and territories globally (Chen et al., 2007). This crop belongs to the genus *Gossypium*, which comprises nearly 50 species. Among these, upland cotton (*Gossypium hirsutum* L.) stands out as the principal source of natural fiber due to its wide adaptability and higher yield, accounting for approximately 95% of global cotton production (Mangi et al., 2024). However, as the global population continues to grow, the demand for food is increasing, thereby intensifying the competition for limited arable land between cotton and cereal crops (Ali et al., 2019). Furthermore, global warming leads to increasingly unpredictable climatic conditions, adversely affecting cotton cultivation (Zhao et al., 2023). Consequently, breeders have developed early-maturing varieties to reduce the cotton fertility period in response to land competition and the frequency of natural disasters.

Flowering time is a crucial factor influencing the early maturity of cotton. Previous studies have demonstrated that flowering plants achieve precise regulation of their flowering time by sensing both external environmental signals (such as photoperiod and temperature changes) and endogenous signals (including the biological clock, endogenous plant hormones, and age) (Susila et al., 2021; Woloszynska et al., 2019). Among the various environmental signals, seasonal variations in photoperiod are relatively stable and can be predicted by the plant’s biological clock, making it an important cue for regulating plant growth and flowering time (Batista et al., 2024). Photoperiod refers to the duration of daylight and darkness, specifically the alternation between day and night (Cai et al., 2024). The mechanisms by which the photoperiodic pathway regulates flowering have been extensively studied. Plants sense the duration of sunlight through photoreceptor proteins that control the stability of the CONSTANS (*CO*) transcription factor, which subsequently activates the expression of the flower-forming gene FLOWERING LOCUS T (*FT*).

This gene is then translocated to the apical meristematic tissue of the shoot, where it interacts with FLOWERING LOCUS D (*FD*), inducing the expression of genes in the floral meristematic tissue, ultimately triggering the initiation of flower development (Yang et al., 2024). Plants can be categorized into three groups based on their varying responses to photoperiod: long-day plants, short-day plants, and day-neutral plants. Notably, several key genes involved in the photoperiodic flowering pathway, such as *GI*, *phyA*, and *ELF3*, exhibit opposing roles in different plant types (Fraser et al., 2021; Kolmos et al., 2011; Li et al., 2023). Cotton originated in the tropics, with most wild cotton species classified as short-day plants. Early studies indicated that *G. hirsutum*, *G. herbaceum*, and *G. arboreum* exhibited delayed flowering under long-day conditions, while long days simultaneously promoted increased plant height and dry matter mass across most species (Bhatt et al., 1982). Subsequently, during human domestication, upland cotton was gradually adapted to different dimensions of photoperiod and thus proved to be a photoperiod-insensitive day-neutral plant (Zhao et al., 2023). However, a recent study revealed that cotton grown under long-day conditions (16-hour light/8-hour dark cycle) flowered 7-11 days earlier than when cultivated under short-day conditions (10-hour light/14-hour dark cycle), indicating that flowering time in cotton remains influenced by day length (Pan et al., 2024). It is important to note that this article focused solely on the impact of day length on flowering time and did not explore how day length regulates nutrient growth in cotton or the interactions between nutrient growth and flowering. Consequently, further investigation into the photoperiodic regulation of cotton flowering is warranted.

In addition to flowering, plants can also regulate nutrient growth by sensing the length of daylight through the length of time photosynthesis and its metabolites are present (Batista et al., 2024). Studies have shown that Arabidopsis thaliana, a long-day model plant, grows faster under long-day (Wang et al., 2024a). Cannabis is a short-day crop, and a longer photoperiod increases its photosynthetic capacity, which in turn promotes cannabis growth but delays flowering (Šrajer et al., 2022). However, for short-day plants, long-day treatment not only delayed the flowering of guar but also inhibited its nutritional growth (Li et al., 2023). This suggests that the regulatory effects of photoperiod on the nutrient growth of different plants are also different. In the growth cycle of cotton, there is an alternating pattern between nutrient growth and reproductive growth, which are closely interdependent. Nutrient growth supplies the essential materials required for reproductive growth, while reproductive growth, in turn, influences the process of nutrient growth. Therefore, while investigating the mechanism of photoperiod regulation on plant flowering, its effect on nutrient growth and the interaction between nutrient growth and flowering after photoperiod induction should not be neglected.

In this study, we investigated the effects of photoperiod on cotton growth and development by integrating physiological indicators with transcriptomic analysis. We analyzed the expression of growth and flowering-related genes in cotton under different photoperiodic treatments. This research contributes to a deeper understanding of the molecular mechanisms underlying the photoperiodic regulation of cotton growth and development.

## 2. Material and methods

### 2.1. Plant materials and treatment

Pot experiment materials consisted of two cotton varieties with different genotypes, XJ12-2 and XJ15-6, which were provided by the Cotton Research Institute of Hunan Agricultural University. Between the two, XJ12-2 is relatively earlier-maturing than XJ15-6. Full and uniformly sized seeds were soaked in water for 12 h, drained, and sown in plastic pots with a soil weight of 4 kg per pot. The soil in the pots was a 1:1 mixture of substrate and field soil, in which the contents of organic matter, total nitrogen, total phosphorus, and total potassium were 89.13 g/kg, 3.16 g/kg, 1.02 g/kg, and 13.44 g/kg, respectively; and the contents of alkaline dissolved nitrogen, quick-acting phosphorus, and quick-acting potassium were 203.01 mg/kg, 75.12 mg/kg, and 13.44 g/kg, respectively. 203.01 mg/kg, 75.12 mg/kg, and 236.35 mg/kg, respectively; and pH 5.18. Eight pots of each variety were sown in a total of 16 pots, and after sowing, they were randomly divided into two groups to receive two types of photoperiodic treatments: (1) long-day treatment (14-h of light/10-h of darkness) and (2) short-day treatment (10-h of light/14-h of darkness). Except for the light time, the rest of the cultural conditions were kept the same.

### 2.2. Plant growth parameters, photosynthetic parameters, chlorophyll content

Cotton squaring and flowering times were recorded and cotton growth parameters were investigated 45 d after treatment (all cotton was squared) by measuring cotton plant height (distance from cotyledonary node to the tip of the main stem) with a straightedge (scale 1 mm) and the length and width of the largest leaf at the top of the cotton to calculate maximum leaf area (length * width * 0.78). Three cotton plants per treatment were selected to separate the top spread leaves and immediately frozen in liquid nitrogen, after which they were stored at -80L for subsequent transcriptional analysis. Cotton was divided by cotyledon node into aboveground and root parts and dried at 105L for 30 min, followed by drying at 80L until constant weight. The remaining cotton was continued to be cultured until all of them flowered.

Net photosynthetic rate (Pn), stomatal conductance (Gs), intercellular CO_2_ concentration (Ci), and transpiration rate (Tr) were measured in the apical-maximum leaves of cotton using a portable photosynthesis measurement system (Li-6800, LiCor, Lincoln, NE, USA), with the light quantum flux density set at 1000 μmol m^-2^ s^-1^. Relative chlorophyll content was measured using a SPAD-502 chlorophyll meter (SPAD-502 plus, Minolta CameraCo., Ltd., Japan), and each treatment was repeated six times.

### 2.3. RNA isolation, library preparation, and RNA-seq

RNA was extracted from 12 cotton leaf samples using TRIzol reagent (Invitrogen, Carlsbad, CA, USA) according to the manufacturer’s protocol. First, genomic DNA contamination was removed using DNaseI (TaKara). The integrity and quality of the RNA were subsequently tested using a 2100 Bioanalyzer (Agilent Technologies, Inc., CA, USA) and an ND-2000 (NanoDrop Technologies, Thermo Fisher Scientific, Madison, USA). High-quality RNA transcriptome libraries were prepared using a TruSeqTM RNA sample preparation kit (Illumina, Inc., SanDiego, California, USA). RNA-Seq was performed on the Illumina NovaSeq 6000 system of Wuhan Mytel Biotechnology Co., Ltd. (Wuhan, China). Raw measurements were cleaned using fastp (https://github.com/OpenGene/fastp) to obtain high-quality data. The purified reads were then aligned to the cotton genome (*Gossypium hirsutum* L. acc. TM-1, https://mascotton.njau.edu.cn/info/1054/1118.htm) and HISAT. Use NR (https://www.ncbi.nlm.nih.gov/Protein/), GO (http://geneontology.org/), KEGG (https://www.kegg.jp/kegg/ko.html), and other public databases for annotating gene functions. The differential expression between the two groups was analyzed using DESeq2 software (1.20.1).

### 2.4. qRT–PCR validation

Transcriptome results were validated by randomly selecting 7 degrees from RNA-seq data. Transcription levels of these genes were subsequently assessed using qRT-PCR. Primers were designed for selected genes using Primer Premier 6.0. These primers were subsequently used to amplify the target genes (Table S1). qRT-PCR analysis was performed using Talent qPCR Master Mix Kit (Tiangen Biotechnology, China) and Roche LightCycler 480 platform (Roche, Switzerland). The internal reference gene is Vactin. The 2^-ΔΔCt^ method is used to calculate the relative expression of genes at the transcription level.

### 2.5. Statistical analysis

One-way ANOVA was used for statistical analysis of the data, SPSS 23.0 was used for least significance difference (LSD) analysis, and the significance level was P < 0.05. The result is the mean ± standard error of 3 replicates. Graphs were drawn using GraphPad Prism 10.0.

## 3. Results

### 3.1 Plant phenotype and photosynthesis

With treatment time, XJ12 under LD treatment was the first to flower, followed by LDXJ21-11, at which time, the two varieties under SD treatment were still in square (Fig.1A, B). The statistical results showed that the squaring time of XJ12 under the LD condition was 35 d, which was 6 d earlier than that under the SD condition. The squaring time of XJ21-11 under LD condition was 39 d, which was 5 d earlier than that under SD condition. The flowering time of XJ12 and XJ21-11 under LD conditions was 12 d and 13 d earlier than that under SD, respectively (Fig.1C). In addition, LD also promoted the growth of cotton plant height and biomass. 22.87%, 82.35%, and 21.06% increases in plant height, root dry weight, and aboveground dry weight were significantly observed in XJ12 under LD conditions compared with SD conditions, while 21.13%, 48.39%, and 15.08% increases in plant height, root dry weight, and aboveground dry weight were observed in XJ21-11 compared with SD conditions. In addition, the maximum leaf area of XJ12 and XJ21-11 decreased by 17.01% and 17.14% under LD compared to SD (Fig. 1D, E).

**Fig. 1.**
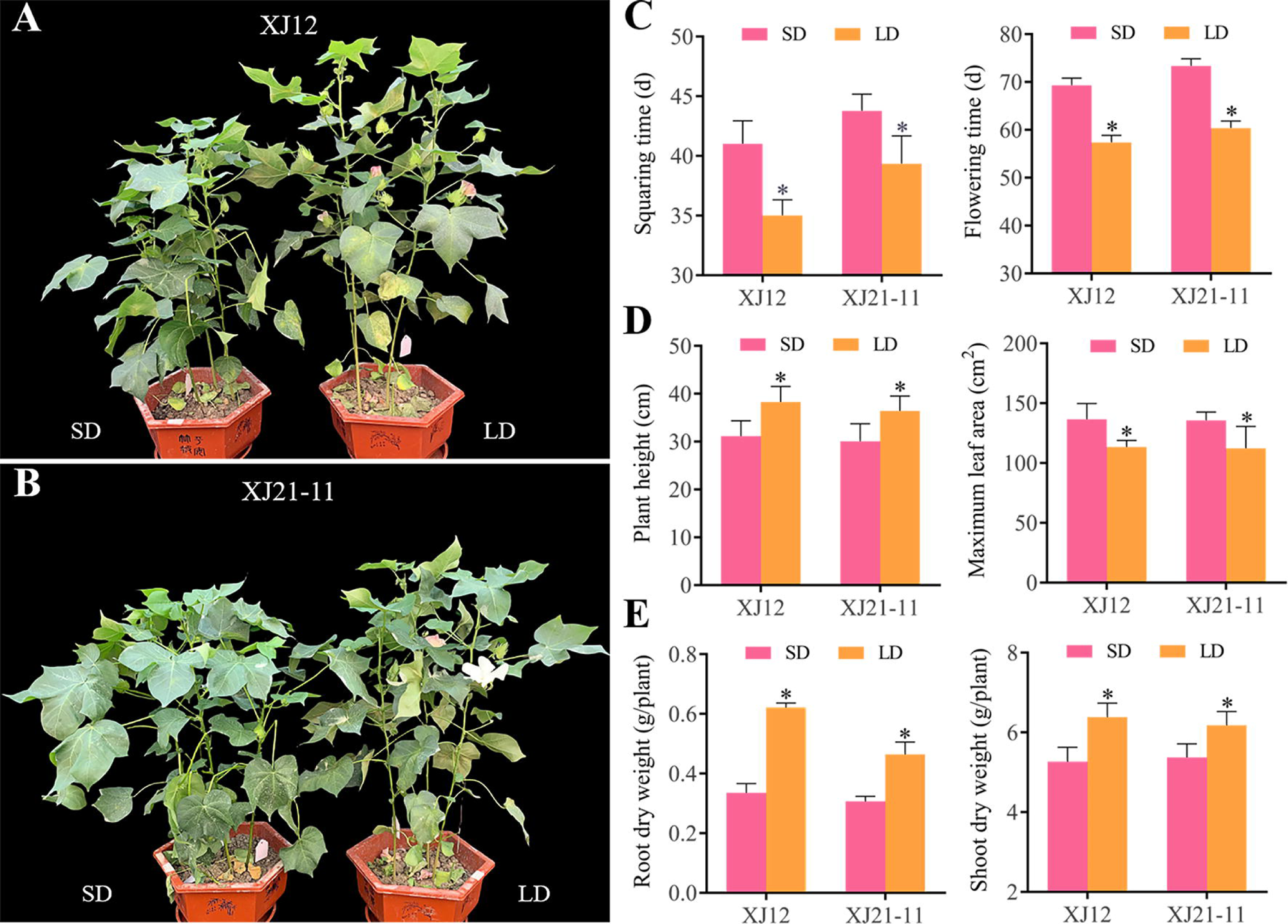
Effect of different photoperiods on cotton phenotype. (A-B) Comparison of cotton flowering under different photoperiod treatments (taken on the 60th day after treatment). (C) Squaring time and flowering time. (D) Plant height and maximum leaf area. (E) Root dry weight and shoot dry weight. Results represent mean values ± SD. One-way ANOVA was used to compare the significant differences of treatments. The statistical significance was defined as * P < 0.05.

We also observed that LD treatment enhanced photosynthesis and SPAD value in cotton (Table 1). Compared with SD, Pn, Gs, Ci, and Tr were increased by 154.79, 240.00, 10.27 and 217.06%, respectively, in XJ12 and 111.86, 171.43, 10.74 and 155.56%, in XJ21-11 under LD treatment. LD treatment significantly increased XJ12 and XJ21-11’s SPAD value were significantly increased by 25.44% and 22.75%, respectively.

**Table 1.** Effect of different photoperiods on photosynthetic parameters and SPAD value in cotton.

| Treatment | Pn<br>($\mu\text{mol}/\text{m}^2/\text{s}$) | Gs<br>($\text{mol}/\text{m}^2/\text{s}$) | Ci<br>($\mu\text{mol}/\text{mol}$) | Tr<br>($\text{mmol}/\text{m}^2/\text{s}$) | SPAD |
| --- | --- | --- | --- | --- | --- |
| LDXJ12 | 17.02±1.45a | 0.34±0.03a | 309.40±14.68a | 5.39±0.54a | 44.77±1.99a |
| SDXJ12 | 6.68±1.30c | 0.10±0.02c | 280.59±18.42bc | 1.70±0.32c | 35.69±2.02c |
| LDXJ21-11 | 11.61±1.79b | 0.19±0.05b | 292.25±17.17ab | 3.22±0.76b | 42.09±2.37b |
| SDXJ21-11 | 5.48±1.82c | 0.07±0.03c | 263.91±14.50c | 1.26±0.48c | 34.29±1.85c |

These values represent the mean values ± SD. Different lowercase letters in the same column indicate a statistically significant difference between treatments ( $p < 0.05$ ).

### 3.2. Transcriptional analysis at different photoperiods

RNA sequencing was conducted on 12 samples (Four treatments were applied, each with three replicates), yielding a total of 47.85-56.73 million reads (Table S2). Of these, 46.84-55.51 million reads were mapped to the reference genome (*Gossypium hirsutum* L. acc. TM-1), resulting in a mapping rate of 97.76% to 97.92%. After filtration, the Q30 values for all samples exceeded 94.51%, and the GC content ranged from 44.85% to 44.98%. Principal component analysis revealed that the 12 samples were distinctly categorized into four groups based on variety and treatment, with the three biological replicates of each sample clustering together (Fig. 2A). Pearson correlation analysis further indicated a high degree of agreement in the RNA sequencing data among the three biological replicates (Fig. 2B). Collectively, these results demonstrate the high reliability of the RNA-seq data and their appropriateness for subsequent bioinformatics analysis. The criteria for screening differentially expressed genes (DEGs) in this study were |log2Fold Change| ≥ 1 and FDR < 0.05.

**Fig. 2.**
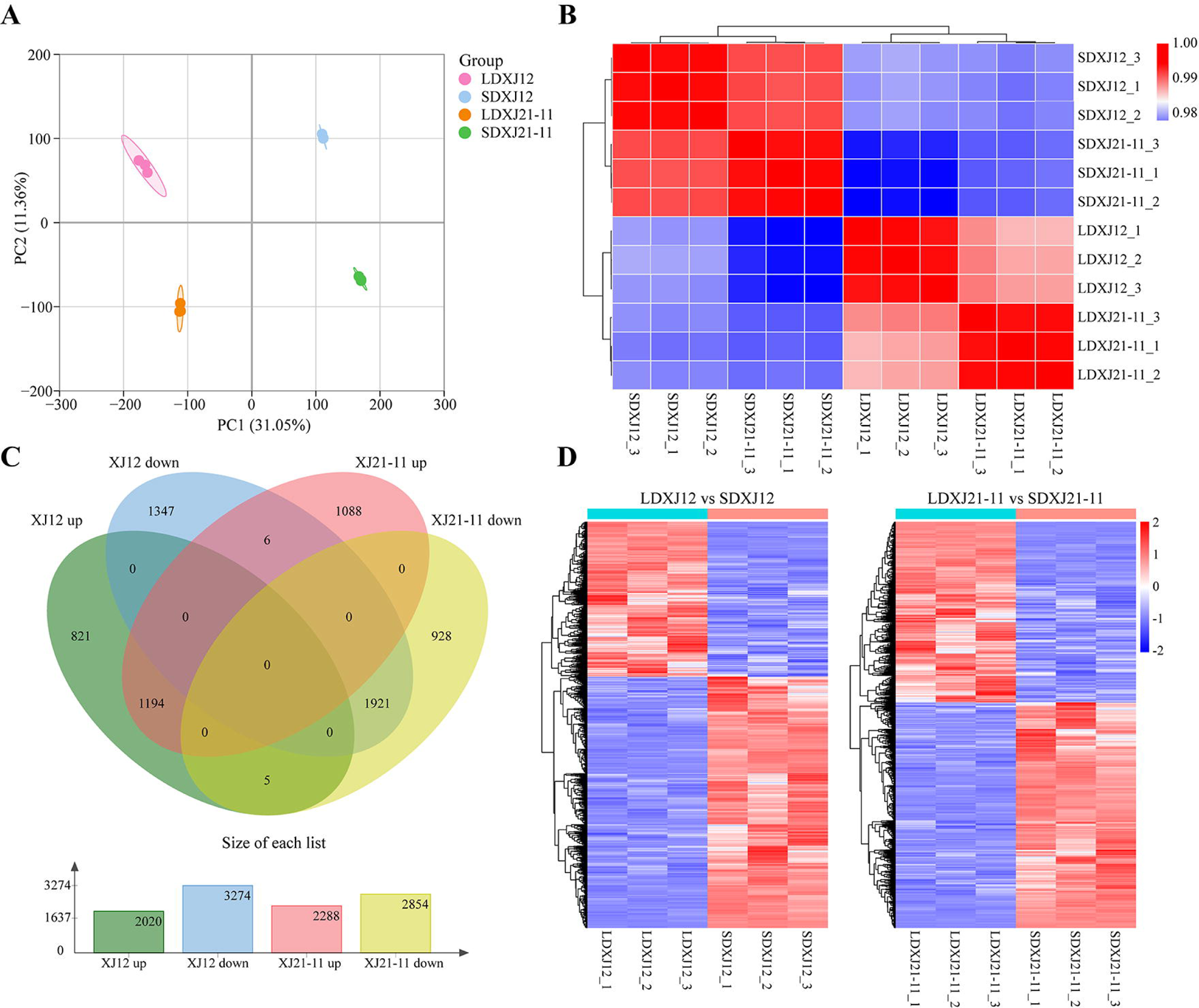
Display of transcriptome data and DEGs under different photoperiods. (A) Principal component analysis of transcriptome data. (B) Correlation analysis of samples. (C) Venn diagram of DEGs. (D) Heatmap of DEGs.

Compared to the SD group, 5,294 DEGs were identified in XJ12 (2,020 up-regulated and 3,274 down-regulated), while 5,142 DEGs were identified in XJ21-11 (2,288 up-regulated and 2,854 down-regulated). Among these, 1,194 DEGs were up-regulated in both cultivars, and 1,921 DEGs were down-regulated in both cultivars (Fig. 2C). This finding indicates that the expression profiles of the two varieties at the transcriptome level are highly similar under different photoperiod treatments. The heat map shows the differences in expression patterns of all DEGs between the two varieties under LD and SD treatments (Fig. 2D).

### 3.3. GO and KEGG enrichment analysis of DEGs

To further elucidate the transcriptional response of cotton to photoperiod, we conducted Gene Ontology (GO) and Kyoto Encyclopedia of Genes and Genomes (KEGG) pathway annotation analyses on the differentially expressed genes (DEGs). The top 30 significantly enriched GO terms are presented in Fig. 3A and 3B. In the XJ12 group, we identified 19 biological processes (BPs), 1 cellular component (CC), and 10 molecular functions (MFs), whereas the XJ21-11 group comprised 15 BPs, 4 CCs, and 11 MFs. Notably, 17 terms are significantly enriched together in both varieties. The two most prominent terms in BP for both varieties were “photosynthetic electron transport in photosystem II” and “photosynthesis, light reaction.” The shared CC term between the two varieties was “photosystem,” while the top three MF terms included “electron transfer activity” and “chlorophyll binding” for both varieties. The top 20 KEGG enrichment pathways are illustrated in Fig. 3C, D, revealing 13 common enrichment pathways across the two cultivars. These pathways predominantly encompassed “Photosynthesis,” “MAPK signaling pathway-plant,” “Plant hormone signal transduction,” “Biosynthesis of secondary metabolites,” and “Metabolic pathways.” These results indicate that the mechanisms underlying photoperiod treatment in XJ12 and XJ21-11 exhibit significant similarities, primarily influencing the signaling and metabolic processes in cotton.

**Fig. 3.**
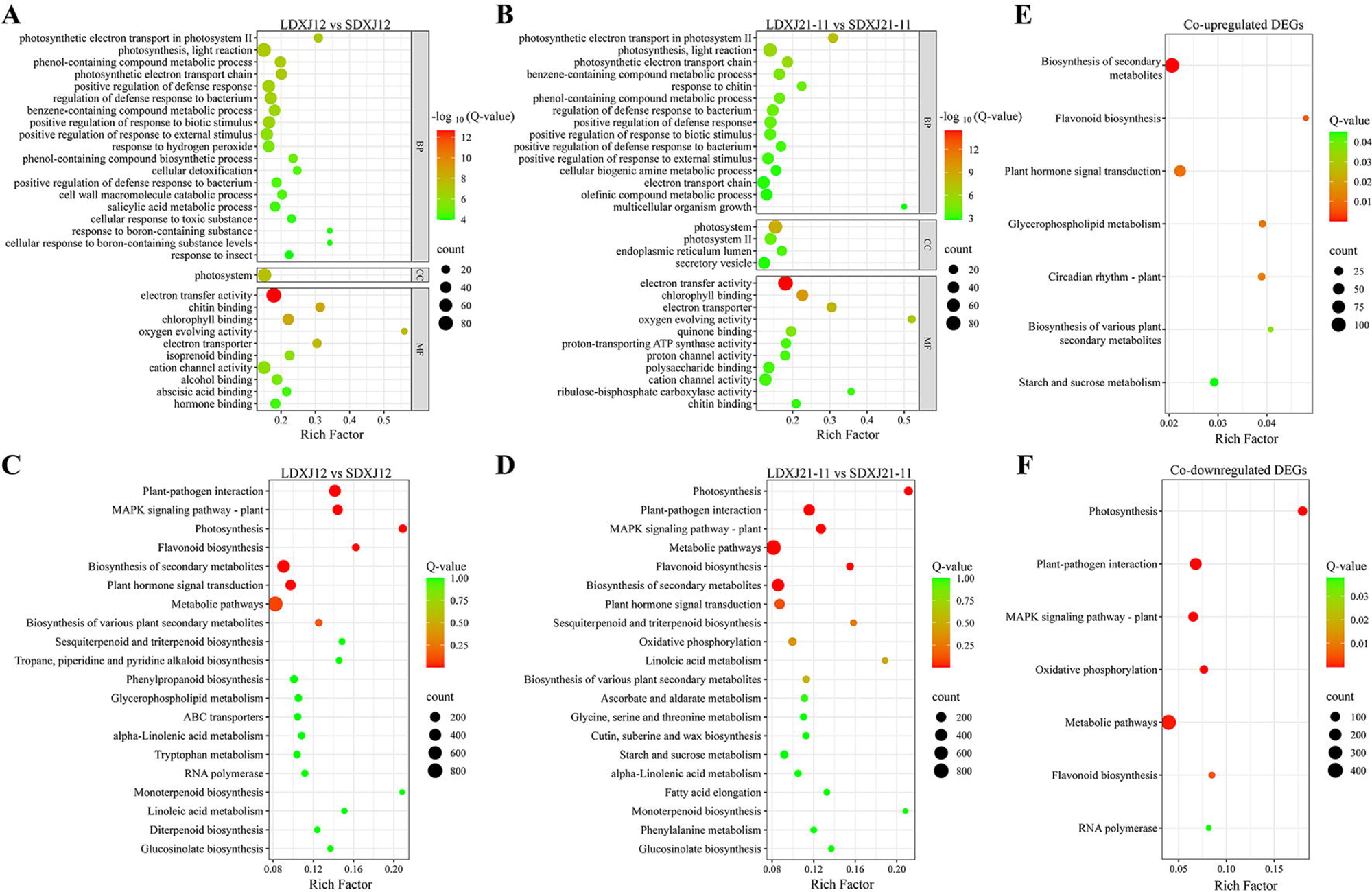
GO and KEGG enrichment analysis of DEGs. (A) Top 30 GO enrichment analysis (LDXJ12 vs SDXJ12). (B) Top 30 GO enrichment analysis (LDXJ21-11 vs SDXJ21-11). (C) Top 20 KEGG enrichment analysis (LDXJ12 vs SDXJ12). (D) Top 20 KEGG enrichment analysis (LDXJ21-11 vs SDXJ21-11). (E) KEGG enrichment analysis for co-upregulation. (E) KEGG enrichment analysis for co-downregulation.

To further investigate the common regulatory mechanisms of photoperiod across different genotypes of cotton, we conducted KEGG enrichment analysis on the differentially expressed genes (DEGs) shared between XJ12 and XJ21-11. The results indicated that the up-regulated DEGs were significantly enriched in the pathways of “Flavonoid biosynthesis,” “Plant hormone signal transduction,” “Circadian rhythm-plant,” and “Starch and sucrose metabolism” (Fig. 3E). Conversely, the down-regulated DEGs were associated with the pathways of “Photosynthesis,” “MAPK signaling pathway-plant,” “Oxidative phosphorylation,” and “Flavonoid biosynthesis” (Fig. 3F).

### 3.4. DEGs involved in photosynthesis

In this study, various photoperiod treatments resulted in differential expression of genes associated with photosynthesis, leading to the selection of 31 differentially expressed genes (DEGs) for further analysis (Fig. 4A). Genes encoding the Light-harvesting chlorophyll protein complex (*Lhcb2*, *Lhcb6*), Photosystem II (*Psbw*), and Photosynthetic electron transport (*PetF*) were found to be up-regulated in the long-day (LD) group compared to the short-day (SD) group. Conversely, a greater number of genes, including those encoding Photosystem II (*PsbA*, *PsbD*, *PsbC*, *PsbB*, *PsbK*, *PsbM*, *PsbH*), Photosystem I (*PsaA*, *PsaB*, *PsaC*), Cytochrome b6/f complex (*PetA*, *PetB*), and F-type ATPase (*alpha*, *beta*, *epsilon*), were down-regulated in the LD group. Additionally, most of the up-regulated genes exhibited higher expression levels in the XJ12 group, whereas the majority of down-regulated genes were more prominently expressed in the XJ21-11 group.

**Fig. 4.**
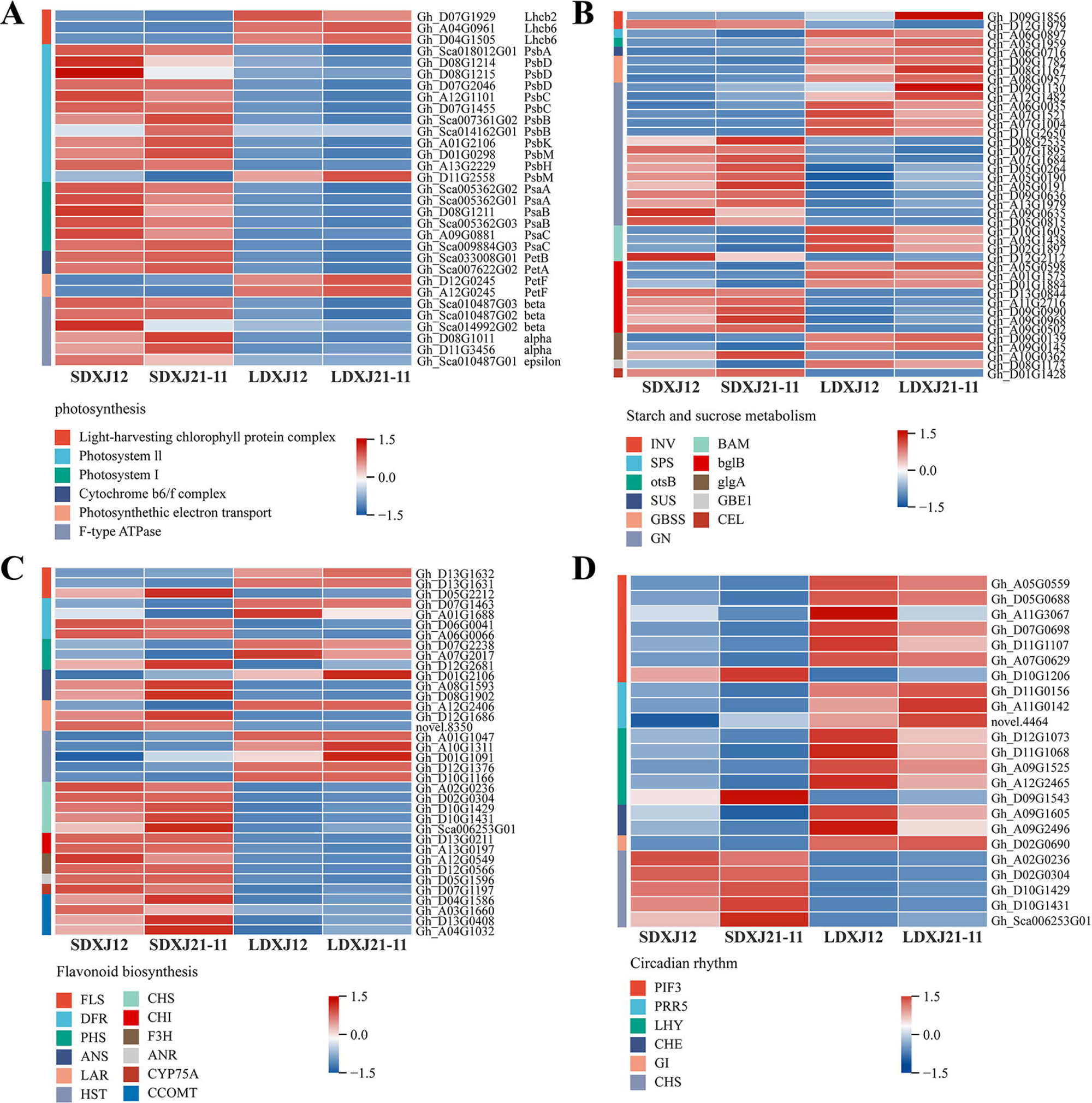
The expression levels of DEGs related to photosynthesis (A), starch and sucrose metabolism (B), flavonoid biosynthesis (C), and circadian rhythm (D) in cotton leaves under different photoperiods.

### 3.5. DEGs involved in starch and sucrose metabolism

Starch and sucrose metabolism are particularly important during plant growth and development, and many DEGs associated with them were identified in our study, including *INV*, *SPS*, *otsB*, *SUS*, *GBSS*, *GN*, *BAM*, *bglB*, *glgA*, *GBE1*, and *CEL*. DEGs encoding *SPS*, *otsB*, *SUS*, *GBSS*, and *GBE1* were up-regulated in the LD group. DEGs encoding *CEL* were down-regulated in the LD group. In addition, DEGs encoding *INV*, *GN*, *BAM*, *bglB*, and *glgA* were also significantly regulated by photoperiod, but their DEGs were either up-regulated or down-regulated in the same treatment group (Fig. 4B).

### 3.6. DEGs involved in flavonoid biosynthesis

KEGG enrichment analysis showed that flavonoid biosynthesis was significantly enriched in both varieties (Fig.3C, D). Flavonoid biosynthesis involves several genes, including *FLS*, *DFR*, *PHS*, *ANS*, *LAR*, *HST*, *CHS*, *CHI*, *F3H*, *ANR*, *CYP75A*, and *CCOMT*. The expression levels of the *HST* gene were significantly up-regulated in the cotton leaves under the long-day treatment compared to the SD group, while *the* expression levels of *the CHS*, *CHI*, *F3H*, *ANR*, *CYP75A*, and *CCOMT genes* were significantly down-regulated. In addition, individual DEGs encoding *FLS*, *DFR*, *PHS*, *ANS*, and *LAR* were either up-regulated or down-regulated in the same treatment group (Fig.4C).

### 3.7. DEGs involved in circadian rhythm

Circadian rhythm-related genes are significantly regulated by photoperiod (Fig. 4D). DEGs encoding *PRR5*, *CHE*, and *GI* genes were up-regulated in the LD group. In addition, multiple DEGs encoding *PIF3* and *LHY* genes were also significantly up-regulated in the LD group compared to the SD group. In contrast, *CHS* genes were down-regulated in the LD group.

### 3.8. DEGs involved in plant hormone signal transduction

Phytohormone signaling plays a role in the central regulatory module of photoperiodic signaling, and we analyzed changes in the expression levels of hormone signaling genes, including Auxin (AUX), Cytokinine (CTK), Gibbere llin (GAs), Ethylene (ET), Salicylic acid (SA), Jasmonic acid (JA), Abscisic acid (ABA) and Brassinosteroid (BR), and most of the DEGs were enriched in AUX, GAs, ABA and BR (Fig.5).

**Fig. 5.**
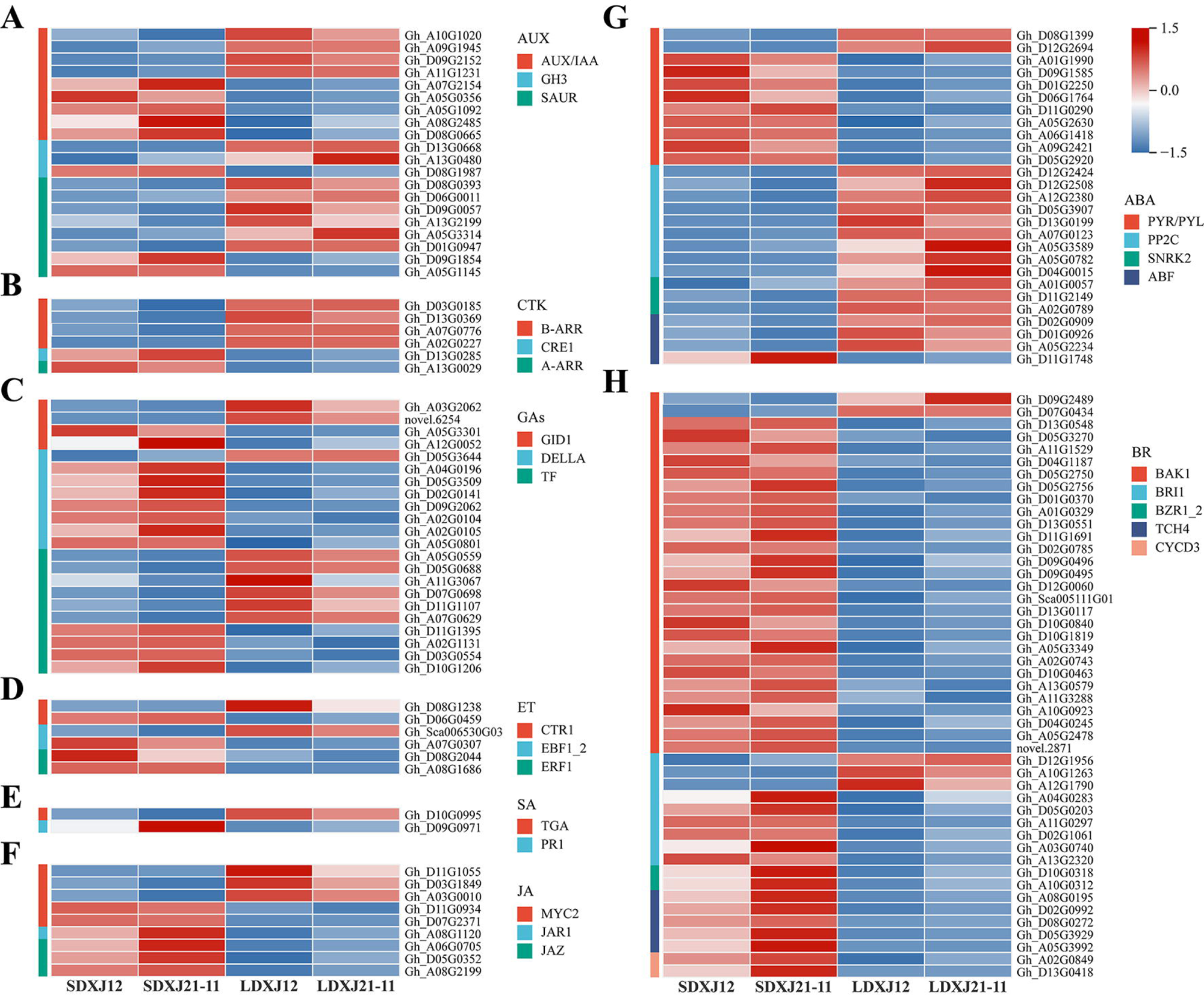
The expression levels of DEGs related to plant hormone signaling in cotton leaves under different photoperiods. (A) Auxin. (B) Cytokinine. (C) Gibbere llin. (D) Ethylene. (E) Salicylic acid. (F) Jasmonic acid. (G) Abscisic acid. (H) Brassinosteroid.

The majority of transcripts in the AUX-related *GH3* and *SAUR* genes were up-regulated in the LD group; a total of eight DEGs were identified in the GAs-related *DELLA* genes, of which seven were down-regulated in the LD group; the majority of transcripts in the ABA-related *PYR/PYL* genes were down-regulated in the LD group, and those in the genes coding for *PP2C* and *SNRK2* were up-regulated in the LD group; and the majority of transcripts in the BR-related *BZR1_2*, *TCH4* and *CYCD3* genes were down-regulated in the LD group, as were the majority of transcripts in the *BAK1* gene; CTK-related *B-ARR* genes were up-regulated in the LD group, and the expression of the *CRE1* and *A-ARR* genes were down-regulated; ET-related *ERF1* genes were down-regulated in the LD group; and *JAR1* and *JAZ* genes related to JA were down-regulated in the LD group; only 2 SA-associated DEGs were identified in our study, of which the *TGA* gene was up-regulated in the LD group, and the *PR1* gene in contrast.

In addition, photoperiodic treatment also significantly regulated AUX-related *AUX/IAA* genes, GAs-related *GID1* and *TF* genes, ABA-related *ABF* genes, BR-related *BRI1* genes, ET-related *CTR1* and *EBF1_2* genes, and JA-related *MYC2* genes, but individual transcripts of the above-mentioned genes were up-regulated or down-regulated in terms of expression in the same group.

### 3.9. DEGs involved in MAPK signaling pathway

The MAPK pathway is responsible for transducing external signals into the cell interior, thereby triggering a series of physiological responses. Our study showed that photoperiodic treatment significantly enriched genes associated with the MAPK signaling pathway, including *FLS2*, *BAK1*, *WRKY22*, *CTR1*, *EBF1_2*, *MYC2*, *PYR/PYL*, *PP2C*, *SNRK2*, *CAT1*, *ER/ERLs*, *MKS1*, *WRKY33*, *MPK3/6*, *VIP1*, *PR1*, *ACS6*, *ERF1*, *CHIB*, *MAPKKK17_18*, *MAPK1_2*, *CALM*, *RBOHD*, and *YODA* genes.

Encoding *PP2C*, *SNRK2*, and *CAT1* were upregulated in long-day treatment. In contrast, genes encoding *MKS1*, *WRKY33*, *MPK3/6*, *VIP1*, *PR1*, *ACS6*, *ERF1*, *CHIB*, *MAPKKK17_18*, *MAPK1_2*, *CALM*, *RBOHD* and *YODA* were down-regulated by long-day treatment. In addition, different transcripts of *FLS2*, *BAK1*, *WRKY22*, *CTR1*, *EBF1_2*, *MYC*2, *PYR/PYL*, and *ER/ERLs* were up-regulated or down-regulated in the same set of expression (Fig. S1A).

### 3.10. DEGs involved in transcription factors (TFs)

Transcriptional regulation plays a crucial role in precisely regulating plant flowering time, and many key flowering-regulated genes belong to the transcription factor family. In our study, *AP2/ERF-ERF*, *MYB*, *WRKY*, *bHLH,* and *NAC* were the most abundant TF families. In addition, genes encoding *MADS-MIKC*, *MADS-M-type,* and *MYB-related* were significantly up-regulated under long-day. Notably, most DEGs encoding *MADS-MIKC* genes were expressed at higher levels in XJ12 than in XJ21-11 (Fig. S1B).

### 3.11. Weighted gene co-expression network analysis (WGCNA)

We conducted a WGCNA analysis of a total of 7,310 differentially expressed genes (DEGs) identified in XJ12 and XJ21-11. Genes with correlation coefficients greater than 0.85, adhering to scale-free network principles, were grouped into a single module, and a soft threshold (power) of 5 was selected. A clustering dendrogram was constructed based on the correlation of gene expression (Fig. 6A), resulting in the identification of seven co-expression modules. Module-trait correlation analysis revealed that the turquoise module (r = 0.99, p < 0.01) was positively correlated with flowering time (FT), whereas the blue module (r = -0.98, p < 0.01) exhibited a negative correlation with FT (Fig. 6B). This indicates that the blue module may play a crucial role in promoting early flowering in cotton under long-day conditions, while the genes within the turquoise module may inhibit flowering. Subsequently, these key modules were subjected to further analysis. Gene Ontology (GO) enrichment analysis indicated that the turquoise module included terms such as “photosynthetic membrane,” “chloroplast thylakoid membrane,” “photosynthesis,” and “jasmonic acid-mediated signaling pathway.” In contrast, the blue module was associated with “regulation of DNA-templated transcription,” “response to light stimulus,” “regulation of transmembrane transport,” and “response to red or far-red light” (Fig. 6C, D). Additionally, KEGG enrichment analysis demonstrated that the turquoise module was significantly enriched in pathways related to “Photosynthesis,” “MAPK signaling pathway-plant,” “Flavonoid biosynthesis,” and “Metabolic pathways.” Conversely, the blue module showed significant enrichment in “Plant hormone signal transduction,” “Biosynthesis of secondary metabolites,” “Circadian rhythm - plant,” and “Starch and sucrose metabolism” (Fig. 6E, F).

**Fig. 6.**
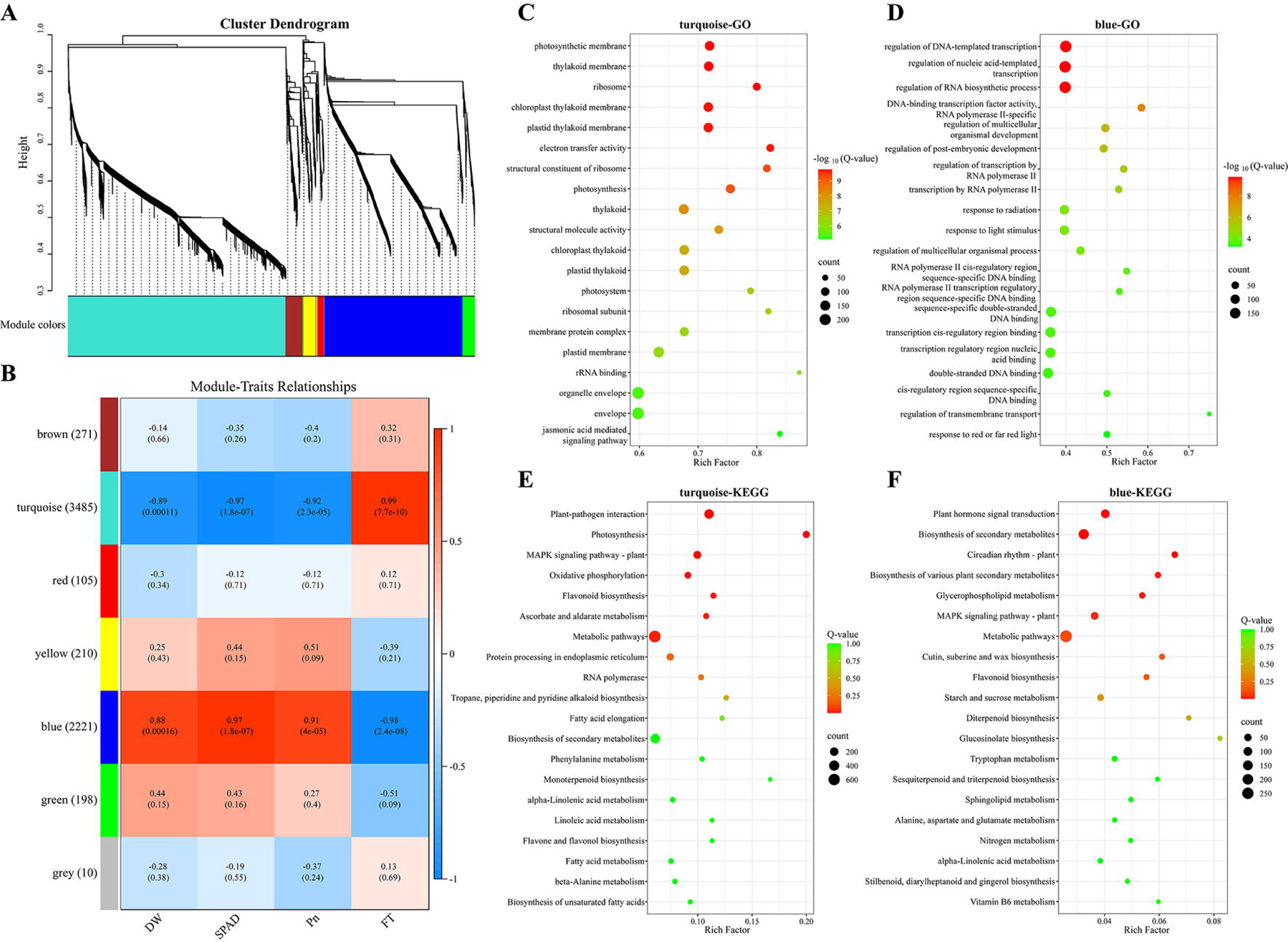
Weighted gene co-expression network analysis of DEGs under different photoperiods. (A) Displays a hierarchical clustering tree for co-expression modules. (B) Correlation heat map of gene co-expression network module and physiological indicators. The numbers within each module represent the correlation value (upper) and p-value (lower). DW is the dry matter mass of cotton per plant; FT is the flowering time. (C-D) Analysis of the top 20 GO enrichment of the turquoise and blue modules. (E-F) Analysis of the top 20 KEGG enrichment of the turquoise and blue modules.

To identify key genes within the modules, we visualized the co-expression networks of the turquoise and blue modules, screening the top 10 central genes in each (Fig. S2). In the turquoise module, we identified one gene encoding brassinosteroid insensitive 1-associated receptor kinase 1 (*BAK1*) (Gh_A01G0329), one gene encoding chalcone synthase (*CHS*) (Gh_A02G0236), one gene encoding mitogen-activated protein kinase 3 (*MPK3*) (Gh_A03G0291), one WRKY transcription factor (Gh_A03G0210), and one NAC transcription factor (Gh_A03G1332). The blue module included one gene encoding mitogen-activated protein kinase 19 (*MPK19*) (Gh_A04G0385), two MADS-MIKC transcription factors (Gh_A03G0634 and Gh_A07G0605), one gene encoding glucan endo-1,3-beta-glucosidase (*GN*) (Gh_A07G1004), and one bHLH transcription factor (Gh_A05G0559). Additionally, we randomly selected seven genes for qRT-PCR analysis. The results indicated that the gene expression patterns determined by qRT-PCR were largely consistent with those observed in the transcriptome analysis (Fig. S3A). Fitting the qRT-PCR data to the transcriptome data yielded an R² of 0.85 (Fig. S3B), demonstrating the accuracy and reliability of the transcriptome data.

## 4. Discussion

It is now widely accepted that upland cotton (*Gossypium hirsutum* L.) has evolved from a perennial, photoperiod-sensitive plant into a day-neutral annual crop as a result of prolonged domestication by humans (McGarry et al., 2016). Our results, however, demonstrated that flowering time was significantly advanced under long-day treatment (14 h light/10 h dark) in both XJ12 and XJ21-11. Additionally, the long-day treatment enhanced the photosynthetic capacity of cotton and promoted its growth (Fig. 1, Table 1). Transcriptome analysis revealed significant differences in gene expression in cotton leaves subjected to different photoperiodic treatments. These findings contribute to a deeper understanding of the molecular mechanisms underlying the photoperiodic regulation of cotton growth and flowering time.

### 4.1. Effect of photoperiod on cotton growth and photosynthesis

Photoperiodic regulation of plant growth primarily occurs through photosynthesis. Extended light durations can enhance the photosynthetic rate, resulting in increased dry weight and biomass of plants (Wang et al., 2024b). Our results further indicate that long-day treatment significantly elevated the photosynthetic rate of cotton, thereby promoting dry matter accumulation (Fig. 1, Table 1). This enhancement is attributed to the up-regulation of the *Lhcb2*, *Lhcb6*, and *PetF* genes under long-day conditions (Fig. 4A). The *Lhcb2* and *Lhcb6* genes are integral to the light-harvesting chlorophyll protein complex, which captures light energy and transfers it to the reaction centers of the photosystem (Lu et al., 2021). *PetF*, a crucial gene in the photosynthetic electron transport chain, encodes ferredoxin, which facilitates NADPH production. This suggests that cotton can more effectively utilize light energy for photosynthesis under long-day conditions, resulting in enhanced growth rates. Notably, in our study, several genes associated with Photosystem II, Photosystem I, the Cytochrome b6/f complex, and F-type ATPase were down-regulated under long-day conditions (Fig. 4A). This down-regulation is attributed to excess light energy, which induces photo-oxidative damage in cotton leaves, leading to the degradation of photosynthetic pigments and negatively impacting photosynthetic efficiency (Dann et al., 2021). Consequently, cotton selectively reduces the expression of these genes following prolonged light exposure to mitigate excessive light energy capture and potential photodamage. Previous studies have demonstrated that extended light exposure accelerates leaf cell division and expansion, resulting in increased leaf size (Xu et al., 2024). However, our study yielded contrasting results, in which the leaf area of the largest leaf of the main stem of cotton was significantly reduced under long-day (Fig. 1). This phenomenon suggests that plants in long-day environments may prioritize an increase in the number of leaf cells as a means of adaptation, rather than merely expanding the size of individual cells.

Sucrose and starch are the primary products of plant photosynthesis. Under adequate light conditions, plants preferentially synthesize sucrose, which is subsequently transported to various locations (Zeng et al., 2021). The synthesis of sucrose predominantly involves key enzymes such as sucrose-phosphate synthase (*SPS*) and sucrose synthase (*SUS*) (Bahaji et al., 2014). The present study demonstrated that the expression of *SPS* and *SUS* was significantly up-regulated under long-day conditions compared to short-day conditions (Fig. 4B), thereby promoting sucrose synthesis and metabolism. Additionally, starch metabolism in plants adapts to changes in photoperiod; under varying photoperiods, plants modulate the rates of starch synthesis and breakdown to ensure an adequate energy supply when photosynthesis ceases at night (Feugier et al., 2013). However, as the photoperiod shortens, plant growth becomes increasingly constrained by the reduction in carbon reserves accumulated during the day (Batista et al., 2024). Consequently, in our study, the starch-related *GBSS* and *GBE1* genes exhibited down-regulated expression under short-day conditions (Fig. 4B) in response to the shortage of photosynthetic products. Furthermore, we observed that in the starch and sucrose metabolic pathways, trehalose 6-phosphate phosphatase (*otsB*) was significantly up-regulated under long-day conditions (Fig. 4B). Previous studies have indicated that *otsB* not only reflects sucrose levels but also responds to day length in leaf slices, generating signals that initiate the flowering program (Wahl et al., 2013). Our results align with these findings, suggesting that long-day conditions promote nutrient growth in cotton and induce flowering by activating the *otsB* gene to release signals.

### 4.2. Effect of photoperiod on flavonoid biosynthesis

Flavonoids are the most prevalent plant secondary metabolites, influencing not only the color of plant flowers but also playing a crucial role in environmental stress responses, protection against UV radiation, mitigation of oxidative stress, and disease resistance (Falcone et al., 2012; Schijlen et al., 2004). Our results indicate that photoperiodic treatment regulates the expression of several genes associated with the flavonoid biosynthesis pathway, including *DFR*, *HST*, *CHS*, *CHI*, *F3H*, and *ANR* (Fig. 4C). Previous studies have demonstrated that at least four key genes involved in flavonoid synthesis (*CHI*, *F3H*, *DFR*, and *UFGT*) are activated in sweet potato leaves under long-day conditions, thereby enhancing the synthesis of flavonoid compounds (Carvalho et al., 2010). Furthermore, extended light exposure can improve photosynthetic efficiency and increase the carbon resources converted into flavonoid compounds (Jaakola et al., 2010). However, one study reported that the knockdown of the *DFR* gene in tobacco diminished energy metabolism and photosynthetic light responses in flowering tobacco, while promoting carbon sequestration during dark responses (Jiang et al., 2023). Interestingly, our findings revealed that long-day treatment significantly down-regulated the expression of several genes related to flavonoid biosynthesis, including *CHS*, *CHI*, *F3H*, and *ANR*, while only the *HST* gene was up-regulated (Fig. 4C). Consequently, we hypothesize that the down-regulation of these genes may be linked to an adaptive mechanism during flowering, wherein the down-regulation allows cotton to allocate more resources toward reproductive growth rather than the synthesis of secondary metabolites like flavonoids. Supporting this hypothesis, studies have shown that the transcripts of *CHS*, *ANS*, and *F3H* genes gradually decrease with flowering during the development of rhododendrons (Nakatsuka et al., 2008).

### 4.3. Effect of photoperiod on circadian rhythm

Photoperiod serves as a reliable signal for plants to perceive seasonal changes, with plants measuring the length of daylight hours through a biological clock located in their leaves. While the circadian rhythm of plants is regulated by an endogenous biological clock, this clock can vary across different environments (Johansson et al., 2015). Research indicates that photoperiod enables plants to optimize the timing of flowering and seed production by modulating the rhythms of their biological clocks (Wang et al., 2024c). In the long-day model organism Arabidopsis thaliana, the biological clock component GIGANTEA (*GI*) plays a crucial role in forming the GI-FKF1 complex, which, upon activation by light, enhances the expression of Constans (*CO*) under long-day conditions, thereby promoting the transcription of FLOWERING LOCUS T (*FT*) (Sawa et al., 2007). Additionally, the CYCLING DOF FACTOR (*CDF*) family gene acts as a transcriptional repressor of *CO*, while *PRR5* directly represses the expression of *CDF* to stabilize *CO* transcription (Nakamichi et al., 2012). Phytochrome-interacting factors (*PIFs*) have been shown to initiate flowering by sensing and integrating photoperiodic signals (Galvāo et al., 2019). However, in the short-day plant rice, *GI* and *PRR* play opposite roles (Takano et al., 2001). A recent study showed that short-day treatment up-regulated *LHY* expression in guar thereby inducing flowering (Li et al., 2023). This study found that long-day conditions up-regulated the expression of *PRR5* and *GI* genes, with DEGs associated with *LHY* also being up-regulated under long-day treatment. These findings suggest that *GI*, *PRR5*, *LHY*, and *PIF3* collectively promote flowering in cotton.

### 4.4. Effect of photoperiod on plant hormone signal transduction and MAPK signaling pathway

In the present study, KEGG annotation of DEGs revealed that signaling pathways were commonly observed in the long and short-day comparisons of XJ12 and XJ21-11. Notably, the plant hormone signal transduction and MAPK signaling pathways ranked among the top 20 pathways for both varieties (Fig. 3C, D). Additionally, weighted gene co-expression network analysis (WGCNA) indicated that the MAPK signaling pathway was enriched in turquoise modules, demonstrating a positive correlation with flowering time (Fig. 6E). Conversely, plant hormone signal transduction was enriched in the blue module and exhibited a negative correlation with flowering time (Fig. 6F). These findings suggest that the photoperiodic regulation of cotton growth and development is intricately linked to signal transduction.

As a signaling molecule, phytohormones are involved in regulating various processes of plant growth and development and are also tightly regulated by external environmental signals such as photoperiod, temperature, and stress (Li et al., 2023). A previous study reported that phytohormones, including GAs, JA, BR, ABA, ET, AUX, and CTK, play a role in the regulation of plant flowering time (Izawa, 2021). GAs, recognized as a key hormone, were the first to be identified as capable of regulating flowering in plants. The *DELLA* protein, an important repressor in the GAs signaling pathway, interacts with the flowering-promoting factor *CO*, inhibiting *CO*’s binding to NF-YB, which consequently leads to a delay in flowering time (Xu et al., 2016). Furthermore, the level of *DELLA* protein varies with the duration of sunlight. Studies have demonstrated that the expression of *DELLA* proteins decreases under long-day conditions, thereby promoting flowering (Phokas et al., 2021). Our findings align with this, as most transcripts encoding *DELLA* proteins were down-regulated under long-day conditions (Fig. 5C), facilitating cotton flowering. It has also been shown that *DELLA* proteins can interact with *JAZ*, a negative regulator of the JA signaling pathway, to alleviate the repression of *MYC2* (Browse and Wallis, 2019). *MYC2* can transcriptionally activate the *PUB22* gene, forming a positive feedback loop that enhances JA signaling (Wu et al., 2024). Our results indicated that long-day conditions down-regulated the expression of *JAZ* genes while up-regulating several transcripts encoding *MYC2* genes (Fig. 5F). This suggests that long-day treatment promotes JA signaling by decreasing the expression level of *DELLA* proteins. Additionally, the *DELLA* protein can bind to *ERF*, forming a feedback regulatory network in response to ET signaling, which in turn inhibits ET signaling (An et al., 2012).

Abscisic acid (ABA) influences plant growth rates by regulating cell expansion and division. In this study, we observed (Fig. 5G) that long-day conditions down-regulated several transcripts encoding *PYR/PRL* genes, which interact with *PP2C* and reduce its inhibitory effect on *SNRK2*. This interaction resulted in a significant up-regulation of *SNRK2* expression and the regulation of various transcription factors through phosphorylation, thereby initiating signaling pathways downstream of ABA and affecting cotton growth and development (Umezawa et al., 2010). Additionally, the signaling genes for auxin (AUX), brassinosteroids (BR), salicylic acid (SA), and cytokinins (CTK) are also significantly influenced by photoperiod (Fig. 5). The Auxin-responsive GH3 gene family (*GH3*) encodes a growth hormone-binding enzyme in Arabidopsis, mediating light-sensitive signaling (Park et al., 2007). FLOWERING LOCUS C (*FLC*) is recognized as a strong repressor of *FT* in Arabidopsis. During BR signaling, the key factor *BZR1* can bind to the promoter of FLOWERING LOCUS D (*FLD*) and repress its expression, which in turn represses *FLC* (Zhang et al., 2013). In our study, most transcripts in the *GH3* gene family were upregulated in the long-day (LD) group, while *BZR1_2* expression was downregulated in the LD group (Fig. 5A, H). Thus, our results suggest that photoperiod induces a complex signaling network among these hormones, which synergistically regulates cotton growth and flowering by integrating the gibberellin (GAs) flowering pathway.

In our study, multiple genes of the MAPK signaling pathway were significantly regulated by photoperiodic treatments (Fig. S1A), which included multiple transcription factors (e.g., *WRKY22* and *WRKY33*, etc.) as well as genes shared with phytohormone signaling pathways (e.g., *BAK1*, *PP2C*, and *SNRK2*, etc.). These results suggest that the MAPK signaling pathway serves as a crucial intersection for phytohormone signaling, enabling the integration of signals from multiple hormones to activate transcription factors (Fig. S1B), thereby regulating the levels of individual hormones, resulting in accelerated growth and early flowering in cotton under long-day conditions (Fig. 1A).

## 5. Conclusions

Our study elucidates the molecular mechanism by which long-day conditions promote cotton growth and early flowering from a transcriptomic perspective. Extended light duration provides more abundant light energy for cotton growth and enables cotton to photosynthesize for a longer period. Long-day conditions enhance cotton growth by down-regulating photosynthesis-related genes to sustain a higher photosynthetic rate while up-regulating genes are associated with sucrose and starch metabolism. Additionally, circadian and signaling genes facilitate earlier flowering times in cotton by responding to signals related to day length. Our findings also indicate that long-day conditions down-regulate flavonoid biosynthesis-related genes, thereby modifying cotton’s secondary metabolism. Overall, long-day conditions activate transcription factors through various signaling pathways, including phytohormone signaling and the MAPK signaling pathway, which subsequently triggers the expression of numerous functional genes. This cascade enhances the photosynthetic capacity of cotton leaves, improves the synthesis and metabolism of sucrose and starch, and ultimately promotes both growth and the advancement of early flowering in cotton (Fig. 7). These results provide new insights into the molecular mechanisms underlying the effects of photoperiod on cotton growth and development.

**Fig. 7.**
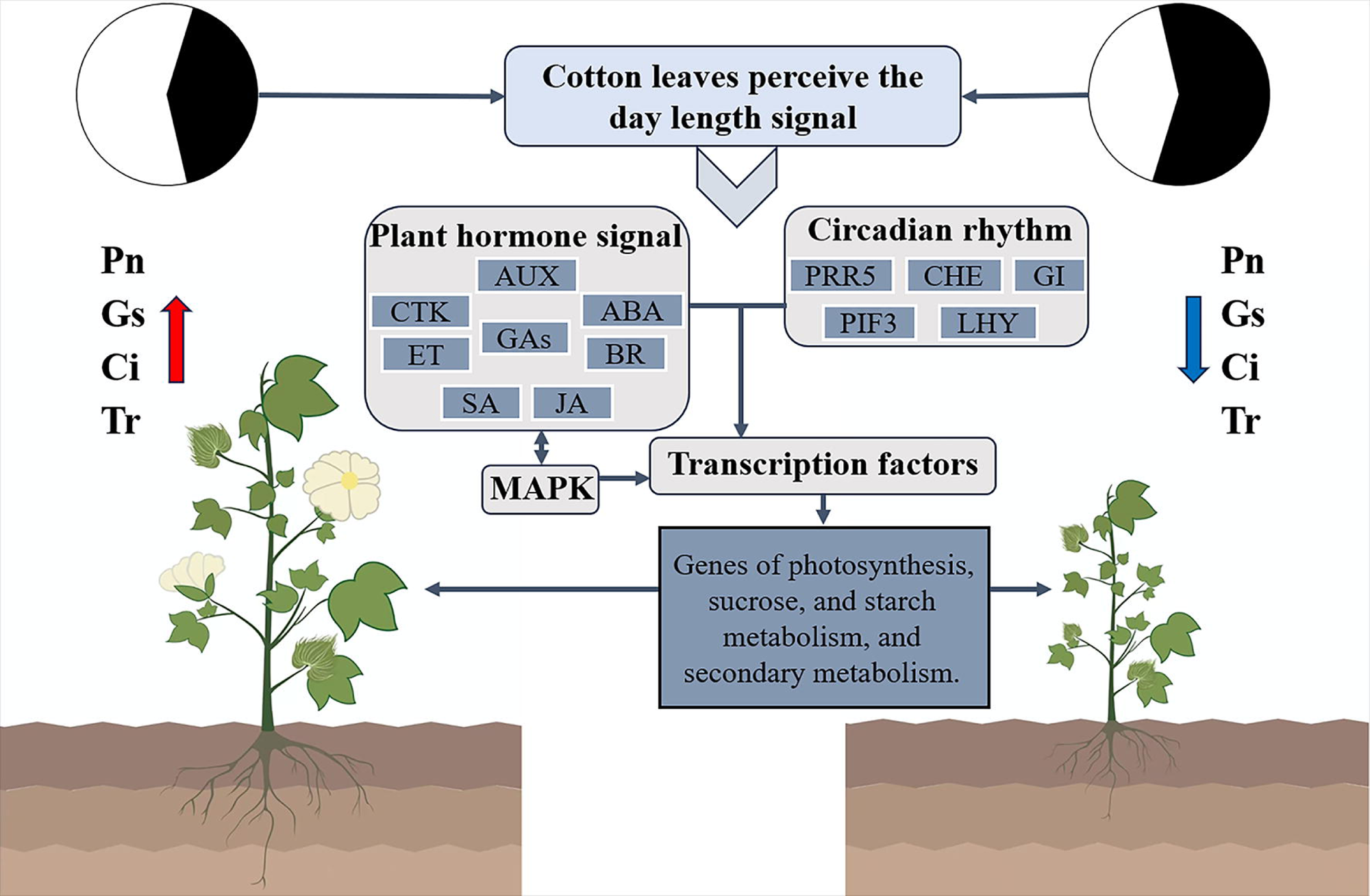
Schematic diagram of the response mechanism of cotton under different photoperiods.

## CRediT authorship contribution statement

**Ning Zhang:** Writing – original draft, Formal analysis, Data curation, Conceptualization. **Yujie Liu:** Writing – original draft, Formal analysis, Data curation. **Haitao Dai:** Methodology, Investigation, Formal analysis. **Penghui Yi:** Validation, Methodology. **Weiwei Cheng:** Validation, Formal analysis. **Xinyu Zhan:** Validation, Formal analysis. **Aiyu Liu:** Conceptualization, Supervision, Writing - review & editing. **Xiaoju Tu:** Writing – review & editing, Supervision, Resources, Project administration, Funding acquisition, Conceptualization.

## Supplementary Figure and Table legends

**Figure S1.** The expression levels of DEGs related to MAPK signaling pathway (A), and transcription factors (B) in cotton leaves under different photoperiods.

**Figure S2.** Correlation network analysis of the top 10 hub genes in the key modules of WGCNA. The shade of color represents the magnitude of connectivity within the module.

**Figure S3.** qRT-PCR validation. The data shown is the mean ± standard error.

**Table S1.** Primer used in qRT-PCR validation.

**Table S2.** RNA-seq data comparison information statistics.

